# Blockade of Kinin B_1_ receptor counteracts the depressive-like behavior and mechanical allodynia in ovariectomized mice

**DOI:** 10.1101/2020.09.01.278416

**Authors:** Izaque de Souza Maciel, Vanessa Machado Azevedo, Patricia Oliboni, Maria Martha Campos

## Abstract

Menopause is related to a decline in ovarian estrogen production, affecting the perception of the somatosensory stimulus, changing the immune-inflammatory systems, and triggering depressive symptoms. Inhibition of kinin B_1_ and B_2_ receptors (B_1_R and B_2_R) inhibits the depressive-like behavior and mechanical allodynia induced by immune-inflammatory mediators in mice. However, there is no evidence on the role of kinin receptors in depressive-like and nociceptive behavior in female mice submitted to bilateral ovariectomy. This study shows that ovariectomized mice (OVX) developed time-related mechanical allodynia and increased immobility time in the tail suspension test (TST). The genetic deletion of B_1_R, or the pharmacological blockade by selective kinin B1R antagonist R-715 (acute, i.p), reduced the increase of immobility time and mechanical allodynia induced by ovariectomy. Neither genetic deletion nor pharmacological inhibition of B_2_R (HOE 140, i.p) prevented the behavioral changes elicited by OVX. Our data suggested a particular modulation of kinin B_1_R in the nociceptive and depressive-like behavior in ovariectomized mice. Selective inhibition of the B_1_R receptor may be a new pharmacological target for treating pain and depression symptoms in women on the perimenopause/menopause period.

## Introduction

In women, the aging process linked to the onset of perimenopause/menopause period, which occurs at approximately 45-55 years of age, is related to a progressive decrease in the circulating levels of gonadal hormones [1–3].

The hormonal changes observed in menopause include an increase in the serum levels of the follicle-stimulating hormone (FSH), associated with diminished estradiol and inhibin B [3]. The decrease in estrogen is associated with several physical and psychological symptoms [2,4–6]. Parts of these symptoms, such as vasomotor alterations, sleep disturbances, depression, sexual dysfunctions, cognitive impairment, and joint pain are specially observed during the perimenopause period. Alternatively, a range of alterations will accompany women for the rest of their lives [7]. Hormone replacement therapy (HRT) remains the most effective therapeutic option to treat menopausal symptoms. However, HRT carries some risks, such as postmenopausal breast cancer, thrombosis, an increase of cardiovascular disease, and stroke [3,8,9].

Besides the importance of estrogen receptors (ER) in mediating the sexual growth and differentiation in women, these receptors are widespread in the central nervous system (CNS) [10]. The activation of the ER in CNS regulates mood, alertness, cerebral blood flow, and neurotransmitter activity [2]. Previous clinical evidence has shown that chronic pain is one of the most prevalent symptoms during perimenopause/menopause [3,11,12]. Furthermore, ER regulates the immune systems and inflammatory mediators [13,14].

Major depression (MD) disorder affects more than 350 million people globally, and the prevalence is higher in women than in men. Noteworthy, data from clinical studies indicated that perimenopause is a critical period for the development of major depression symptoms [4]. Also, the currently available antidepressant therapy showed a mixed result about the effectiveness of treating the MD symptoms. Additionally, many patients do not continue the therapy, because of the variable side effects of antidepressants [15]. Preclinical studies have shown that ovariectomized rats or mice develop depressive-like behavior and nociception changes, which was counteracted by estradiol treatment [16–18]. However, HRT’s adverse effects cannot be disregarded [8]. Thus, the development of new approaches to treating major depression and pain symptoms associated with menopause are required.

During the last years, the kinin receptors have been associated with a series of pathophysiological conditions [19–22]. The biological effects of kinin are mediated via activation of two metabotropic receptors, called B_1_R and B_2_R. While B_2_R is constitutively expressed in several tissues and preferably activated by bradykinin (BK), the B_1_R is weakly expressed under physiological condition but is upregulate after traumatic and immune-inflammatory conditions [23,24]. The activation of B_1_R is modulated by kinin metabolites and contributes to chronic allodynia and hyperalgesia associated with the production of inflammatory cytokines [25,26]. Also, the activation of B_1_R has been implicated in some central nervous system (CNS) conditions, such as Alzheimer’s disease, epilepsy, and bipolar disorder [27–29]. On top of that, our previous work showed that the B_1_R participates in the depressive-like behavior induced by the systemic administration of *E. coli* lipopolysaccharide (LPS) in mice [20]. Such an effect seems to be related to an increase of TNFα levels and microglia activity in the mice brain [20].

Considering the impact of ovarian estrogen depletion on MD, pain, and immune-inflammatory system [4,14], it seems plausible that kinin receptors may participate in behavior changes induced by ovariectomy in rodents [18]. We hypothesized here that inhibition of kinin receptors might attenuate the behavioral changes in ovariectomized mice. Thus, we assessed the pharmacological and genetic inhibition of B_1_R and B_2_R on the mechanical nociception and depressive-like behavior in ovariectomized female mice.

## 2. Materials and Methods

### 2.1. Animals

Experiments were conducted using adult female mice (8 weeks of age with 25 to 30 g), swiss, C57BL6 wild type (WT), B_1,_ and B_2_ receptor knockout mice (B_1_ -/- and B_2_ -/-, UNIFESP-EPM). Animals were housed under standard conditions of light, temperature, and humidity (12 h light-dark cycle, 22 ± 1 °C, under 60 to 80 % humidity), with food and water provided *ad libitum*. Swiss and C57/BL6 mice were obtained from the Central animal facility from the Universidade Federal de Pelotas (UFPEL, Brazil). B_1_ and B_2_ receptor knockout mice were obtained from the animal facility from the Department of Biophysics, Universidade Federal de São Paulo (UNIFESP-EPM, São Paulo, Brazil). All procedures were conducted in accordance with the Brazilian Council for Animal Experimentation (COBEA) guidelines, which comply with international laws and policies for the investigation of experimental pain in conscious animals. The protocols were approved by the local Ethical Committee (protocol number CEUA-PUCRS 12/00290), and all efforts were made to minimize animal suffering and reduce the number of animals used. The experiments were conducted between 08:00-17:00 with randomization of the experimental groups throughout the day.

### 2.2. Ovariectomy surgical procedure

The surgical procedure was conducted as previously described in the literature [16,17], with minor modifications. Briefly, the animals were anesthetized by intraperitoneal (i.p.) injection of ketamine (50 mg/kg) and xylazine (5mg/kg), and after the onset of anesthesia, the lumbar dorsal was shaved. The exposed skin was prepared for aseptic surgery with povidone-iodine (10%) scrub followed by a sterile saline wipe. The ovary was resected bilaterally with skin opened 1- to 2 cm incision in the midline on the lumbar vertebral line, and the ovary was pulled through a small opening in the musculature. A ligature was placed around the exposed ovary and initial segment of the fallopian tube and removed (OVX group). Then the skin and muscle incision were sutured (4-0 nonabsorbable). The sham group underwent the same procedure as the OVX group but without resection of the ovaries and initial segment of the fallopian tube. The behavioral tests were initiated after a recovered period of 7 days.

### 2.3. Measurement of uterine weight

Uterine atrophy is an indicator of the success of the ovariectomy surgical model [16]. After the behavior test, all animals of OVX and Sham group were euthanized by deep inhalation of isoflurane. The uterus of each animal was resected quickly without the periovarian fat and weighted immediately. The values were expressed in mg, and as indicative of success on ovariectomy procedure, the OVX group showed a decrease of uterine weight compared with the Sham group.

### 2.4. Mechanical allodynia

The measurement of the mechanical paw withdrawal threshold (PWT) was carried out using the up-down paradigm, as described in the literature [30,31], with minor modifications. The mice were individually acclimated for one hour in elevated clear plexiglass boxes with a wire mesh floor to allow access to the right hind paws’ plantar surface. Von Frey filaments of increasing stiffness (0.02–10g) were applied to the right hind paw plantar surface of the animals with pressure high enough to bend the filament. Tests were initiated with 0.4g filament. The absence of a paw lifting after 5 s led to the use of the next filament with increased weight, whereas paw lifting indicated a positive response and led to the use of the next weaker filament. This paradigm continued for a total of 6 measurements, including the one before the first paw-lifting response had been made, or until four consecutives positive (assigned a score of 0.030) or four successive negatives (assigned a score of 6.76) responses occurred. The mechanical paw withdrawal threshold response was then calculated as described previously [32], using the following formula: threshold 50% = log of the last hair used − (K. mean); where K is the constant based on Dixon Table, and mean refers to the mean difference (in log units) between stimuli (for mice 0.44). The paw withdrawal threshold was expressed in grams (g) and was evaluated before (baseline) and 7, 14, 21, and 28 days after surgical procedure. A significant decrease in paw withdrawal threshold compared to baseline values was considered as mechanical allodynia.

### 2.5. Tail Suspension Test

According to the methodology described initially [33], we used the tail-suspension test (TST) to assess the depressive-like behavior. At different time-points following the ovariectomy surgical procedure (7, 14, 21, and 28 days), the animals were suspended 50 cm above the floor using adhesive tape, placed approximately 1 cm from the tip of the tail. The mice were exposed to TST, 30 min after the PWT test. The time during which mice remained immobile was quantified in seconds for 6 min [20].

### 2.6. Open-field locomotor activity

The locomotor activity was assessed in the open-field arena 7, 14, 21, and 28 days after the ovariectomy surgical procedure. The experiments were conducted 30 min after TST in a sound-attenuated room, under low-intensity light. Mice were individually placed in the center of an acrylic box (40 × 60 × 50 cm), with the floor divided into nine squares. The number of squares crossed with the four paws was registered for 6 min [20].

### 2.7. Pharmacological treatment

To assess the involvement of B_1_ and B_2_ receptors on the behavior changes induced by ovariectomy surgery. The animals were pre-treated with selective B_1_ receptor antagonist R-715 (0.5 mg/kg; i.p.) and selective B_2_ receptor antagonist Hoe-140 (50 nmol/kg; i.p.). The positive control group was treated with pregabalin (30 mg/kg; i.p.) or saline (0.9%; i.p.). All treatments were administered 30 min before the behavior test PWT test. The doses and the treatment schedules were determined based on previous literature or pilot experiments [20,26,34].

### 2.8. Drugs and Reagents

The following drugs and reagents were used: B_1_ receptor antagonist R-715 was kindly provided by Dr. Fernand Gobeil (Department of Pharmacology, University of Sherbrooke, Qc, Canada) and B_2_R antagonist Hoe-140 was of the commercial source (Bachen, USA; #4043056). Ketamine (Cristalia), Xylazine (Syntec), 10% povidone-iodine, pregabalin (Lyrica®) was obtained from Pfizer (UK). All drugs were diluted in sterile saline immediately before the injections (NaCl 0.9%) solution.

### 2.10. Statistical analysis

The results are presented as the mean ± SEM. The statistical analysis was performed by one- or two-way analysis of variance (ANOVA) followed by Bonferroni *post hoc* test or Student’s t-test. *P*-values less than 0.05 (*p* < 0.05) were considered significant. All tests were performed using the Prism GraphPad Software (San Diego, USA).

## 3. Results

### 3.1. Ovariectomized mice showed mechanical allodynia and depressive-like behavior

Initially, we characterized the behavior changes induced by the menopause surgical model. The ovariectomized Swiss mice (OVX group) showed mechanical allodynia, characterized by a significant reduction in the right hind PWT (OVX = F (1, 20) = 42.72, p = 0.0001, N = 11 by group; at 7, 14 and 21 days, respectively; figure 1B). However, on the 28^th^ day after the surgical procedure, there was no significant difference between groups (OVX x Sham, P = 0.49; figure 1B). The same group of mice demonstrated a significant increase of immobility time in TST (OVX = F (1, 40) = 31.20, p = 0.001, n = 11 by group; at 14, 21 and 28 days after surgery, Figure 1C). The locomotors activity assessed in the open-field arena was not significantly altered (OVX = F (1, 40) = 0.49, p = 0.48, N = 11 by group; figure 1D). Finally, to confirm the surgical procedure, the uterine weight of the OVX group was significantly lower than the sham group (t = 9.72, df = 20; p = 0.0001, figure 1E). Based on these results, the next experiments were done after 21 days of surgical procedure of ovariectomy.

**Figure 1.**
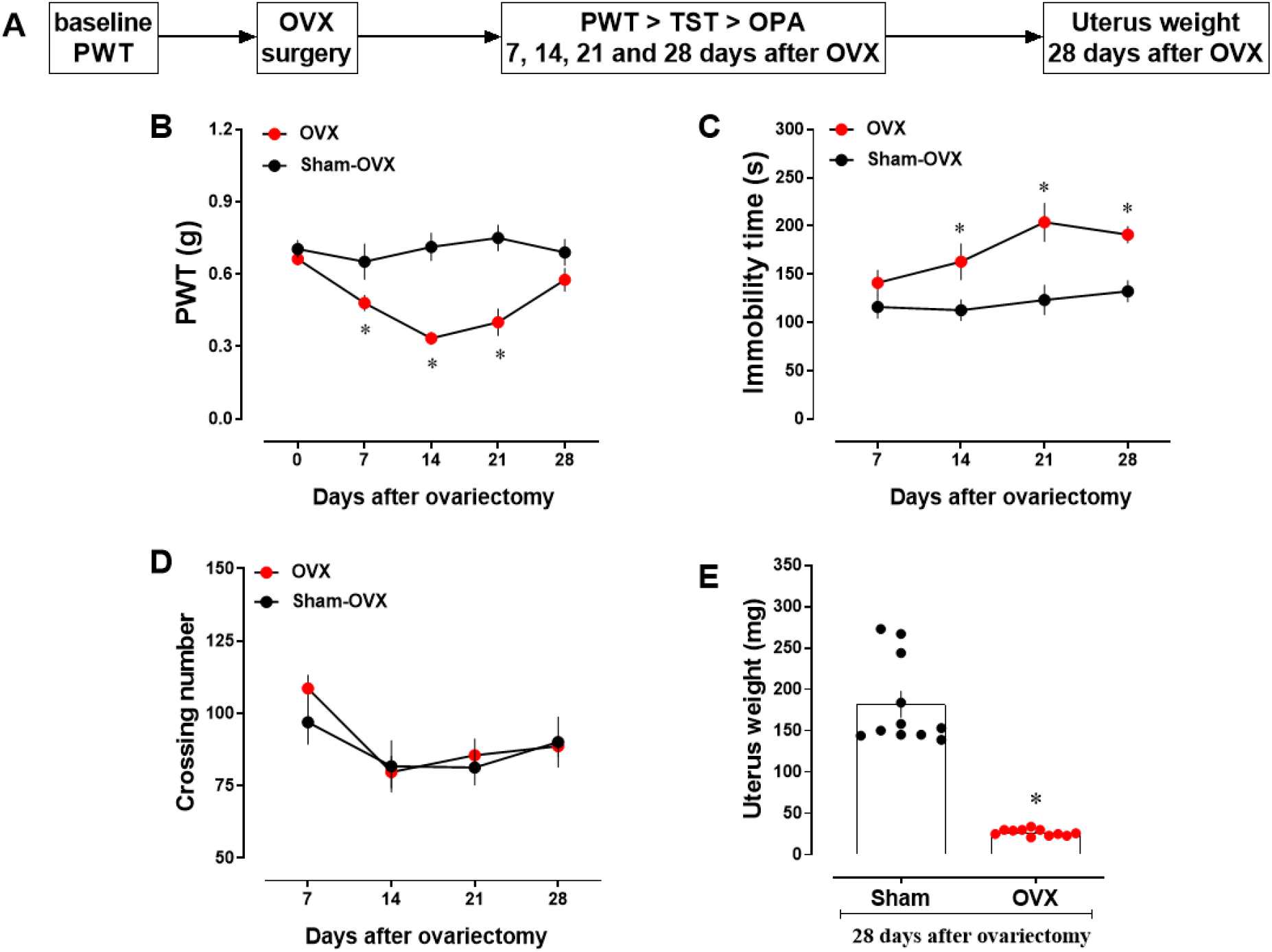
Time-related behavior effect after ovariectomy in mice. (A) Timeline of the experimental approach. Effects of ovariectomy at 7, 14, 21, and 28 days: (B) Mechanical threshold analyzed by using the von Frey test; (C) immobility time in tail suspension test; (D) crossing number in the open-field arena test, and (E) uterus weight (mg) after 28 days of surgery. Each point or column represents the mean ± SEM of 11 animals per group. ^*^P < 0.05 and significantly different from the sham group (two-way repeated-measure ANOVA (time x OVX) followed by Bonferroni post hoc, or Student’s *t-test*). PWT = paw withdrawal threshold; TST = tail suspension test; OPA = open-field arena; OVX = ovariectomy.

### 3.2. B_1_ receptor antagonist R-715 attenuated the mechanical nociceptive and depressive-like behavior in ovariectomized mice

In the next step, we essayed the effects of an effective dose of pregabalin and the B1 receptor antagonist peptide for kinin R-715 in mechanical allodynia (von Frey filaments test) immobility time in the TST test. Our results showed that ovariectomized Swiss mice showed significant decrease in the PWT, when measured at 21 days following the surgical procedure (OVX = F (1, 42) = 29.43, p = 0.0001, N = 13; figure 2B). The acute treatment with pregabalin (30 mg/kg; i.p.; 30 min. before behavior test) or with B_1_R antagonist R-715 (0.5 mg/kg, i.p; 30 min. before behavior test), significantly inhibited the mechanical allodynia induced by ovariectomy (interaction = F (3, 42) = 12.51, p = 0.0001, N = 7-13; figure 2B). However, only o treatment with R-715 significantly reduced the TST immobility time (F (3, 41) = 26,84 p = 0.0001, N = 7-13; figure 2C) in ovariectomized female mice. Both pharmacological treatments were not able to reverse the decrease of uterus weight in ovariectomized mice (p = 0.99, N = 7-13; Figure 2D).

**Figure 2.**
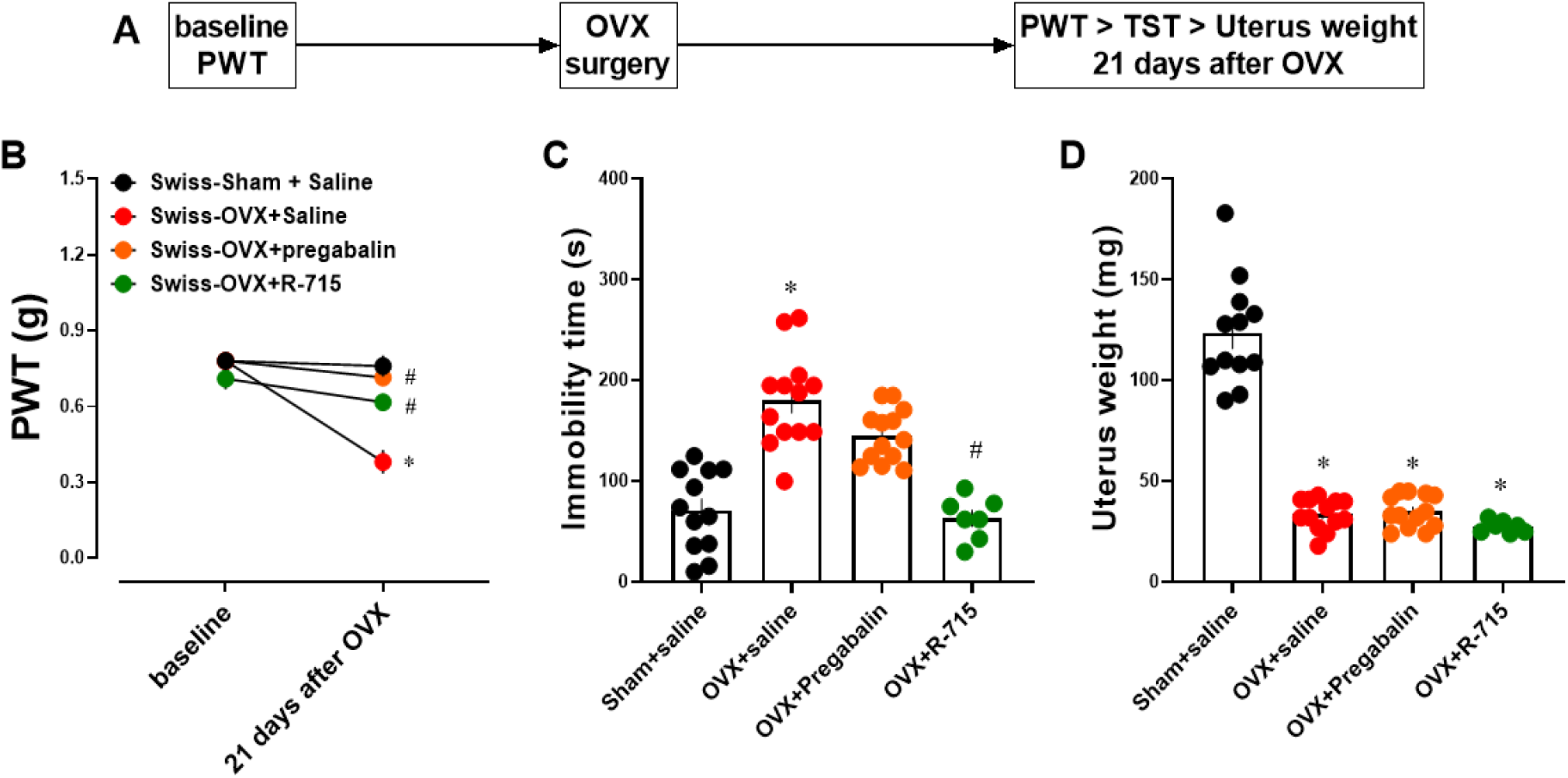
Evaluation of acute treatment with B_1_ antagonist in ovariectomized Swiss mice. (A) Timeline of the experimental approach. Effects of ovariectomy surgical at 21 days. (B) Mechanical threshold analyzed by using the von Frey test, (C) immobility time in tail suspension test, and (D) uterus weight (mg). Effects of treatment with pregabalin (30 mg/kg) or R-715 (0.5 mg/kg); both by i.p., injected 30 min before behavioral tests. Each column represents the mean ± SEM of 7-13 animals per group. ^*^P < 0.05 significantly different from the sham Swiss group; ^#^P < 0.05 significantly different from the OVX Swiss group (one or two-way repeated-measure ANOVA (time x OVX+treatment) followed by Bonferroni post hoc test). PWT = paw withdrawal threshold; TST = tail suspension test; OVX = ovariectomy.

### 3.3. B_1_ receptor knockout mouse is protected of behavior changes elicited by ovariectomy

To confirm the role of the B_1_ receptor for kinin in the mechanical allodynia and depressive-like behavior induced by ovariectomy, we used the B_1_ receptor knockout female mouse (KOB_1_) in our experimental paradigm. Our results showed that ovariectomized C57BL6 mice (WT) showed a significant decrease in PWT when measured at 21 days following the surgical procedure (OVX = F (1, 42) = 14.02, p = 0.0005, N = 10; figure 3B). Both KOB_1_ mice submitted the ovariectomy (OVX-KOB_1_ group) and OVX-WT treated with R-715 (0.5 mg/kg; i.p., 30 min. before behavior test) significantly showed inhibition of mechanical allodynia induced by ovariectomy (interaction = F (4, 42) = 10.19, p = 0.0001, N = 8-10; figure 3B). Relevantly, both OVX-KOB_1_ group and OVX-WT treated with R-715, significantly decreased the immobility time in TST (F (4, 39) = 9.0, p = 0.0001, N = 8-10; figure 3C). Both pharmacological treatments with B1 receptor antagonist or gene deletion were unable to reverse the decrease of uterus weight in mice ovariectomized (p > 0.05; N = 8-10, Figure 3D).

**Figure 3.**
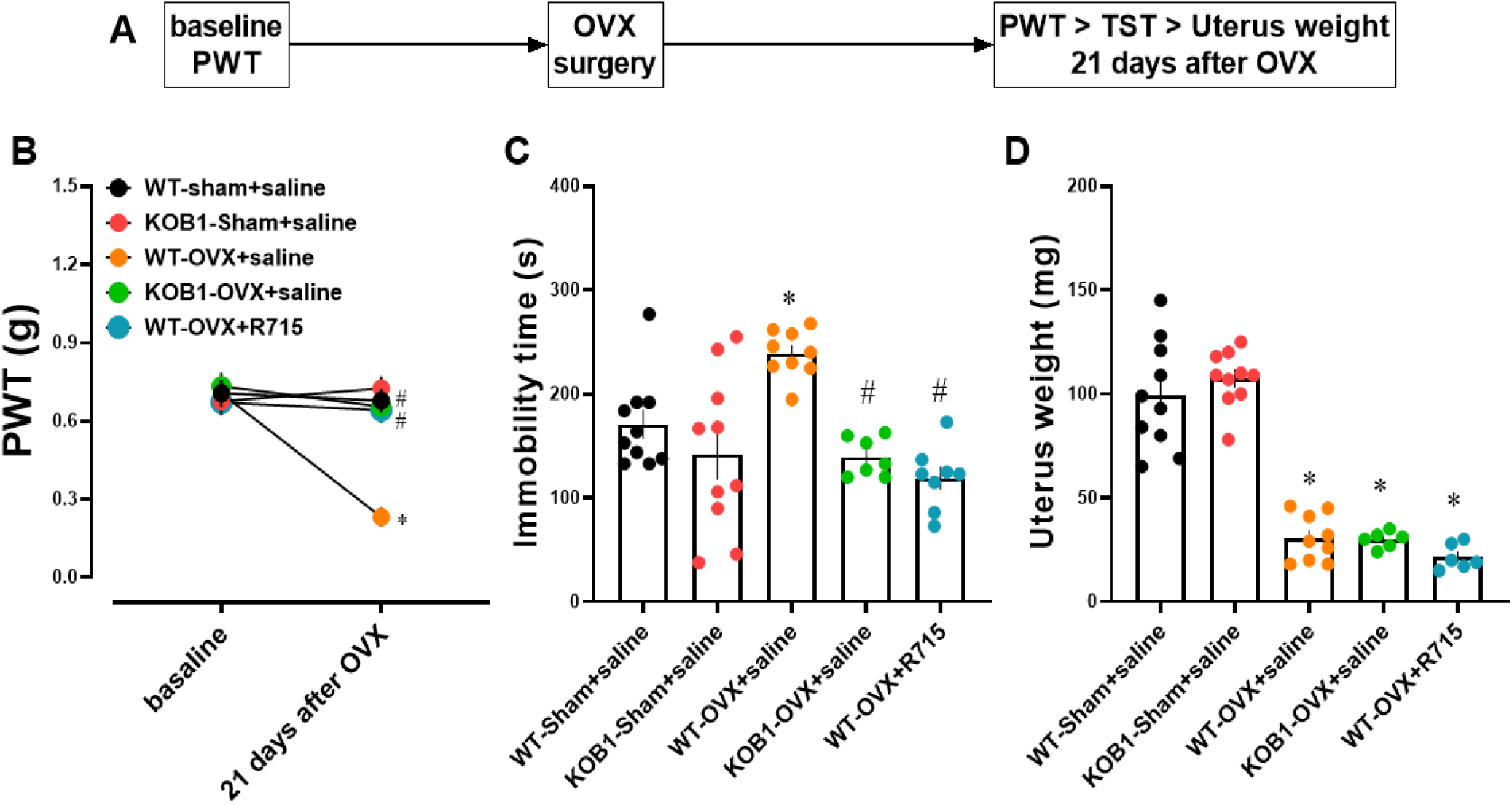
B_1_ receptor showed a protective effect against mechanical allodynia or depressive-like behavior in ovariectomized mice. (A) Timeline of the experimental approach. Effects of ovariectomy surgical at 21 days in C57BL6-WT or KOB_1_ female mice. (B) Mechanical threshold analyzed by using the von Frey test, (C) immobility time in tail suspension test, and (D) uterus weight (mg). Effects of treatment with R-715 (0.5 mg/kg; i.p., 30 min before behavior test). Each column represents the mean ± SEM of 8-10 animals per group. ^*^P < 0.05 significantly different from the sham WT group; ^#^P < 0.05 significantly different from OVX WT group (one or two-way repeated-measure ANOVA (time x OVX+treatment) followed by Bonferroni post hoc test). PWT = paw withdrawal threshold; TST = tail suspension test; OVX = ovariectomy.

### 3.4. Pharmacological inhibition or gene deletion of the B_2_ receptor failed to protect against the behavior changes induced by ovariectomy

We investigated the participation of the B_2_ receptor for kinin in the behavior changes elicited by ovariectomy. We used the B_2_ receptor knockout mouse (KOB_2_) and selective B_2_ receptor antagonist Hoe-140 (50 nmol/kg; i.p, 30 min before behavior test). Similarly, to the OVX-WT group, KOB_2_ female mice that were submitted the ovariectomy (OVX-KOB_2_ group), developed mechanical allodynia when measured at 21 days following the surgical procedure (OVX = F (1, 33) = 129.2, p = 0.0001, N = 6-9; figure 4B). Neither OVX-KOB_2_ group nor OVX-WT group treated with Hoe-140 were not effective in preventing the mechanical allodynia induced by ovariectomy surgery (interaction; F (4, 33) = 17.84, p = 0.99; N = 6-9; figure 4B). Furthermore, both the OVX-KOB_2_ group or OVX-WT treated with Hoe-140, failed to reduce the immobility time in the TST test (F (4, 32) = 18.30, p = 0.99, N = 6-9; Figure 4C). Both pharmacological treatments with B_2_ receptor antagonist or gene deletion were unable to reverse the decrease of uterus weight in mice ovariectomized (p > 0.05, N = 6-9; Figure 4D).

**Figure 4.**
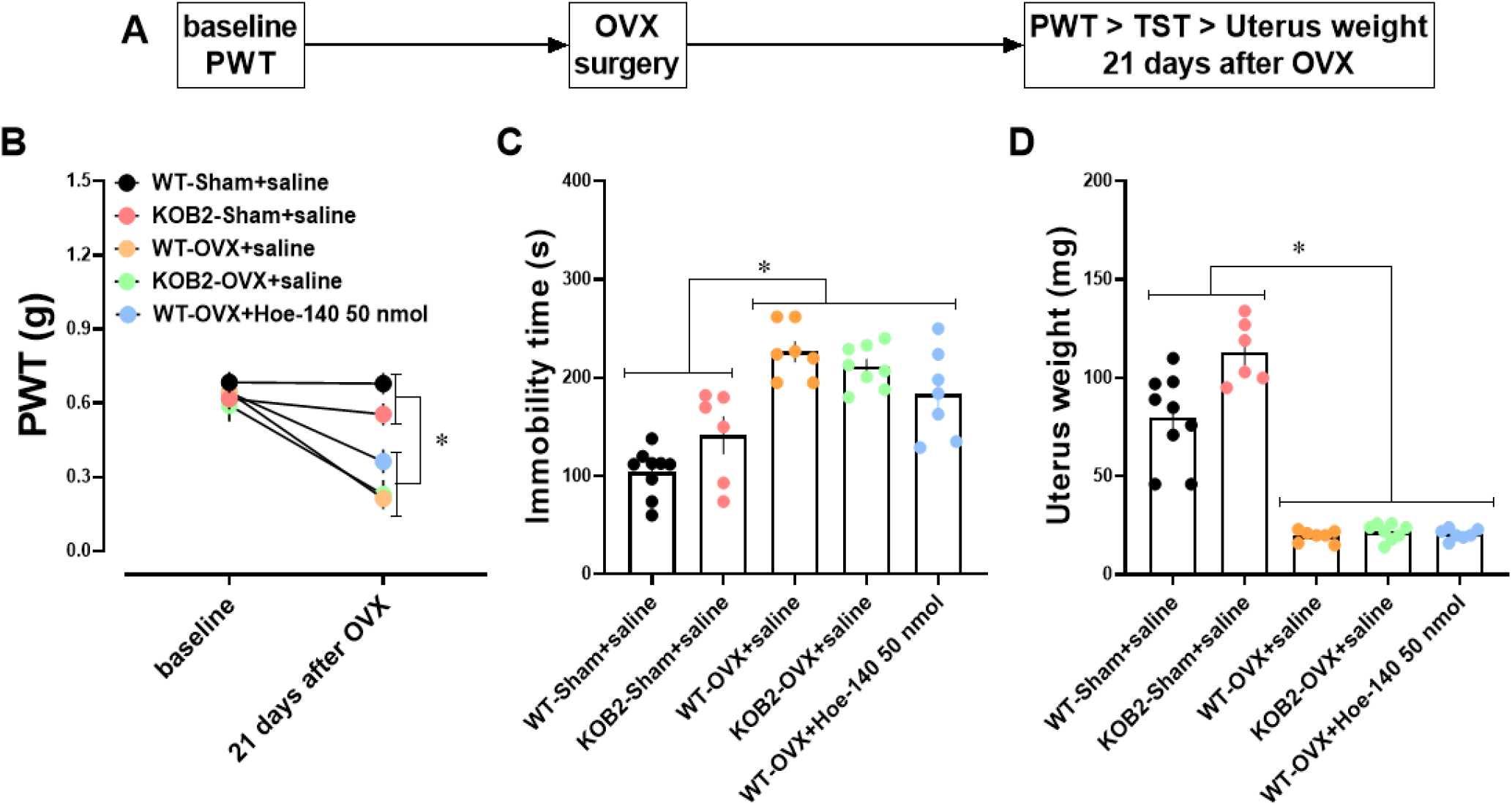
B_2_ receptor failed to inhibit mechanical allodynia or depressive-like behavior in ovariectomized mice. (A) Timeline of the experimental approach. Effects of ovariectomy surgery after 21 days in C57BL6-WT or KOB_2_ mice. (B) Mechanical threshold analyzed by using the von Frey test, (C) immobility time in tail suspension test, and (D) uterus weight (mg). Effects of treatment with Hoe-140 (50 nmol/kg; i.p., 30 min before behavior test). Each column represents the mean ± SEM of 6-9 animals per group. ^*^P < 0.05 significantly different from WT-sham+saline or KOB2-sham+saline group (one or two-way repeated-measure ANOVA (time x OVX+treatment) followed by Bonferroni post hoc test). PWT = paw withdrawal threshold; TST = tail suspension test; OVX = ovariectomy.

## 4. Discussion

The present study describes the participation of bradykinin receptors in the depressive-like behavior and mechanical allodynia triggered by a preclinical model of menopause in mice. Both pharmacological inhibition and genetic deletion of the B_1_ receptor mitigates the behavioral changes induced by ovariectomy. However, neither genetic nor pharmacological inhibition of the B_2_ receptor for kinin attenuated the mechanical hypernociception and increase of immobility time produced by the bilateral ovariectomy. Our data indicate a specificity to the B_1_ receptor subtype for bradykinin in the OVX-induced mechanical allodynia and depressant-like behavior, suggesting a potential new pharmacological target to treat pain and symptoms of major depression in women during the menopause.

During the perimenopause/menopause period, an increase in depression and pain symptoms is associated with a decrease of ovarian hormones [35], and in preclinical studies, the decline of ovarian hormones by bilateral ablation of ovaries induces mechanical allodynia and thermal hyperalgesia and an increase of immobility time in FST or TST [16–18,36]. Corroborating previous work, we observed that bilateral ovariectomy significantly decreased the uterine weight, an indirect indicator of estrogen decrease [16,18]. The uterine atrophy was associated with mechanical allodynia at 7, 14, and 21 days after surgery and a significant increase in the immobility time in the TST at 14, 21, and 28 days. The locomotor activity was not significantly changed compared to the sham group, indicating that an increase of immobility time in TST is not due to motor impairment. Thus, it is plausible to assume that the depletion of ovarian hormone levels facilitates the development of mechanical allodynia and acute coping behavior (helplessness state) in female mice.

Clinical and preclinical studies show the relationship between the decrease of ovarian hormones and increase of pro-inflammatory markers, such as interleukin-1 (IL-1), interleukin-6 (IL-6), and tumor necrosis factor- α (TNF-α) [37,38], which are associated with the molecular mechanisms of pain perception and symptoms of major depression [37,39,40]. It has been postulated that pro-inflammatory cytokines elicited mechanical nociception and acute coping behavior in mice by activating the B_1_ kinin receptor [20,41]. Such mechanisms are associated with an increase in the expression of B_1_R in the peripheral and central structures related to allodynia and major depression [20,24,41–43]. This evidence is in line with the result from the present study, which pregabalin (positive control) and the B_1_R antagonist R-715 effectively inhibited the mechanical allodynia in OVX mice. However, only R-715 was able to decrease the immobility time in TST. On the other hand, the selective B_2_R antagonist HOE-140 did not prevent the increase of mechanical nociception or immobility time in ovariectomized mice. Also, B_1_R, but not B_2_R knockout mice, were unresponsive to OVX-induced mechanical allodynia and increased in the immobility time. Thus, decreasing ovarian hormones in ovariectomized mice may increase B_1_R expression/activity and modulate nociception and depressive-like behavior.

In humans and rodents, the reduction of estrogen followed by ovariectomy is associated with an increase of systemic and central immune-inflammatory mediators [40]. Treatment with 17 beta-estradiol significantly inhibited the increase of microglia activation induced by LPS in the ovariectomized mice [44]. In our previous study, the increase of immobility time in TST induced by systemic injection of LPS in male mice significantly increases the TNF-alpha levels in the whole brain, serum, and cerebrospinal fluid (CSF), followed by an acute increase of B_1_R mRNA expression on the hippocampus and cortex [20].

Besides that, acute systemic treatment with B_1_R antagonists decreased the microglia activation in the hippocampus. Also, TNF-alpha P55 knockout mice treated with LPS do not present depressive-like behavior and increase in B_1_R expression [20]. Together, these data suggest that 17 beta-estradiol and the selective B_1_R antagonist (R-715) may be acting by a similar mechanism to protect against the behavioral changes and molecular alteration modulated by increasing TNF-alpha.

Although still not fully elucidated the molecular mechanism triggered by decreasing of ovarian hormones on the pain process and major depression. Clinical and preclinical data suggest that the decrease of ovarian hormones induce: (I) alteration of angiotensin-converting enzyme (ACE) activity, (II) increase of oxidative stress, and (III) increase of pro-inflammatory cytokines [45–47], which leads to increased production of kinin and receptors activation [40,48].

In the clinical study, the hormone replacement therapy (HRT) in hypertensive postmenopausal women decrease the serum levels of ACE activity and increase the plasma level of angiotensin II (ANG II) and bradykinin [45]. Cultured primary hypothalamic neurons of mice treated with ANG II showed a significant increase of B_1_R expression, an increase of oxidative stress and proinflammatory cytokines, which were prevented by pretreatment with B_1_R antagonist (R-715) [49]. Thus, the HRT may facilitate the kinin-kallikrein system and B_1_R activity.

The reduction of estrogen levels has been associated with increased reactive oxygen species (ROS) [50–52]. In preclinical studies, the increase of ROS induced upregulation of B_1_R on the rat brain submitted to the insulin resistance model [47]. The increase of B_1_R activity leads to downstream pathways involving an increase of nitric oxide (NO), glutamate, and substance P (SP) [47]. Also, ovariectomized mice showed an increase of nitric oxide (NO) in the hippocampus, and the acute treatment with non-selective NO synthase inhibitor (L-NAME) significantly decreases the immobility time of OVX mice in the FST [17]. Thus, decreasing ovarian hormones may increase B_1_R activity via oxidative stress and pro-inflammatory cytokines [17,47].

Together, these data suggest that antagonists of kinin receptors could be a new pharmacological approach to treat pain and major depression symptoms during menopause. However, it is not clear how ovarian hormones modulate B_1_R expression and activity. Further investigation is needed to address the direct role of ovarian hormones on the kinin-kallikrein system related to major depression.

In conclusion, we showed the specificity of B_1_R on mechanical nociception and depressive-like behavior female mice submitted to ovariectomy. To our knowledge, this is the first evidence that the pharmacological or genetic inhibition of the B_1_R prevented the behavioral changes induced by the surgical menopause model in female mice. Thus, the B_1_R inhibition could be a new pharmacological target to treat pain and major depression during the perimenopause/menopause period.

## Conflict of interest and funding

None of the authors declares any conflict of interest. This work was supported by grants from Brazilian research agencies (CAPES), National Council for Scientific and Technological Development (CNPq), Fundação de Amparo à Pesquisa do Estado do Rio Grande do Sul (FAPERGS) and FINEP Research Grant “Implantação, Modernização e Qualificação de Estrutura de Pesquisa da PUCRS” (PUCRS INFRA) # 01.11.0014-00.

## Acknowledgments

The authors thank Dr. Plinio C. Casaratto (University of Helsinki) and M.Sc. Luma Melo (Indiana University) for their review of the manuscript draft.

## Authors’ contribution

ISM designed the study, performed the experiments, collected, analyzed and interpreted data, and wrote the manuscript draft. VMA and PO performed the experiments, collected, analyzed data. MMC designed the study, analyzed and interpreted data, and wrote the manuscript.

## Abbreviations

OVX: ovariectomized mice;
TST: tail suspension test;
PWT: paw withdrawal threshold;
FSH: follicle-stimulating hormone;
HRT: hormone replacement therapy;
ER: estrogen receptors;
MD: major depression;
BK: bradykinin;
CNS: central nervous system;
LPS: lipopolysaccharide;
TNFɑ: tumor necrosis factor-alpha;
CSF: cerebrospinal fluid;
ACE: angiotensin-converting enzyme;
ROS: reactive oxygen species.

## Notes

### Competing Interest Statement

The authors have declared no competing interest.

